# Aggregation Tool for Genomic Concepts (ATGC): A deep learning framework for somatic mutations and other sparse genomic measures

**DOI:** 10.1101/2020.08.05.237206

**Authors:** Jordan Anaya, John-William Sidhom, Faisal Mahmood, Alexander S. Baras

**Author notes:** Corresponding author: Alexander S. Baras^1,2,4^.

## Abstract

Deep learning can extract meaningful features from data given enough training examples. Large-scale genomic data are well suited for this class of machine learning algorithms; however, for many of these data the labels are at the level of the sample instead of at the level of the individual genomic measures. Conventional approaches to this data statically featurise and aggregate the measures separately from prediction. We propose to featurise, aggregate, and predict with a single trainable end-to-end model by turning to attention-based multiple instance learning. This allows for direct modelling of instance importance to sample-level classification in addition to trainable encoding strategies of genomic descriptions, such as mutations. We first demonstrate this approach by successfully solving synthetic tasks conventional approaches fail. Subsequently we applied the approach to somatic variants and achieved best-in-class performance when classifying tumour type or microsatellite status, while simultaneously providing an improved level of model explainability. Our results suggest that this framework could lead to biological insights and improve performance on tasks that aggregate information from sets of genomic data.

---

While deep learning has made significant progress in a range of biological tasks^1^, for genomics data this progress has been limited to predicting features of sequence elements and positions in the genome, such as transcription factor binding, DNAse I sensitivity, and histone-based modifications, or whether the sequence functions as a promoter^2,3^. Making predictions at a higher level, such as at the level of a collection of genomic measures, is complicated by the curse of dimensionality—the high dimensional space makes the data sparse and in general promotes overfitting^4^. Current approaches to this problem include manually reducing the dimensionality through feature selection, dimension reduction techniques such as singular value decomposition, negative matrix factorisation, and various types of autoencoders, or the use of sparse networks which attempt to reduce the weights of the model^5^. However, reducing the dimensions of the data or capacity of the model may produce suboptimal results.

Regardless of how the features of the individual genomic measures are generated, currently a simple aggregation such as a sum or mean is performed to get to a sample-level vector (representing a set of genomic measures). Then, a model such as a random forest or neural net is applied to these sample vectors to perform the sample-level machine learning task at hand (Figure 1). This process essentially weights each genomic measure of the set derived from a given sample equally when in fact it may be that some specific measure(s) are more salient. A more modern attention strategy which dynamically weights genomic measures into sample-level feature vectors may identify these specific measures. Moreover, with current approaches all of the learning occurs at the sample level and “end-to-end” training is not possible, which would allow for novel encoding strategies of genomic measures driven by the machine learning task (Figure 1).

**Figure 1.**
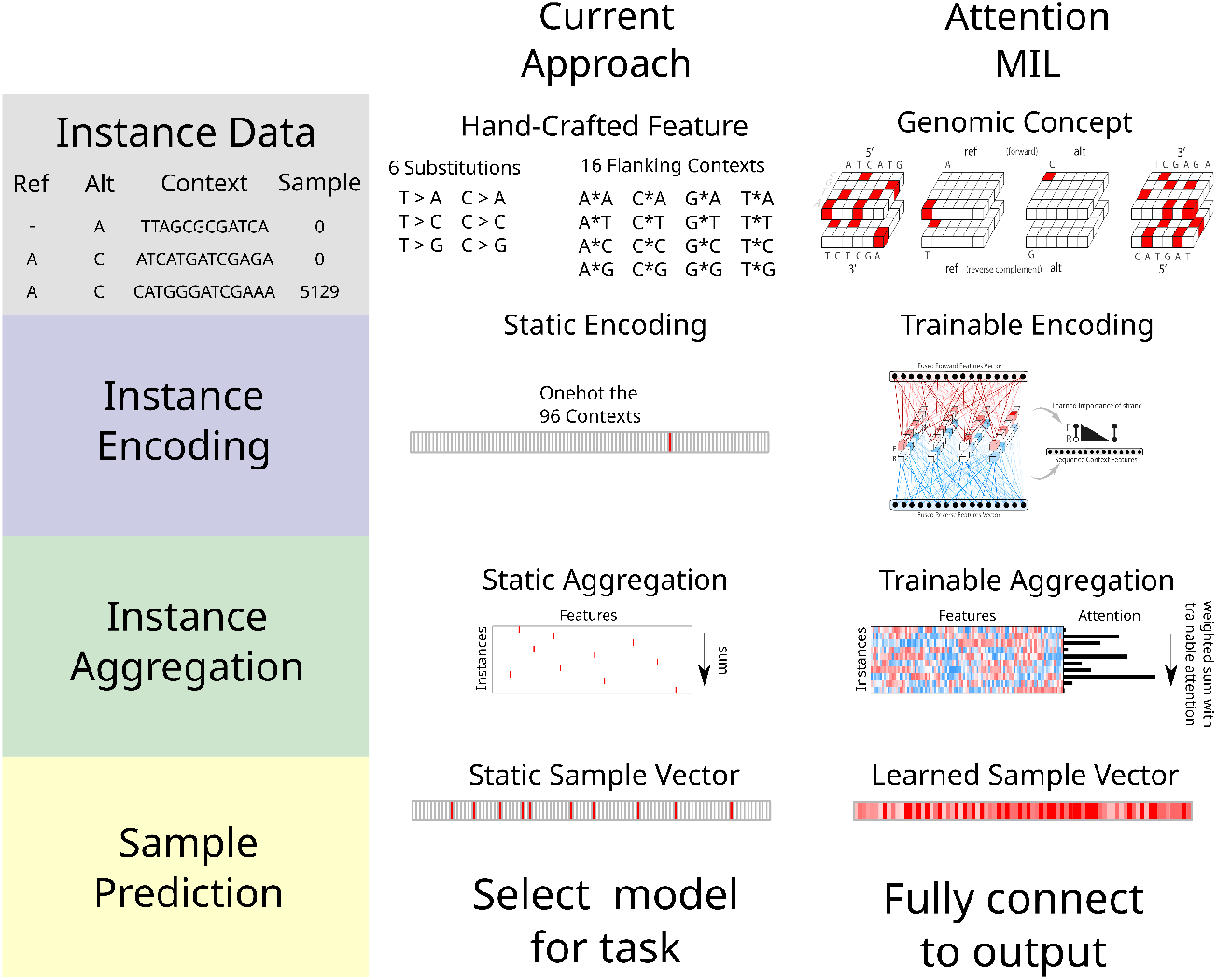
Approaches for sparse genomics data. Data such as somatic mutations must first be encoded and aggregated at the sample level prior to making predictions about a sample (defined as a set of mutations). Currently the process of encoding and aggregating is handled separately from making predictions with the sample-level vectors. With attention MIL it is possible to encode, aggregate, and make predictions with a single end-to-end model. This allows the model to learn a rich feature space at the instance level while also calculating attention for each instance prior to aggregation, thereby allowing for model explainability.

This weakly supervised problem, where features are learned for individual measures (instances) while supervision occurs at the sample level, is the multiple instance learning framework (MIL)^6–8^. MIL has recently revolutionised the field of computational pathology, allowing researchers to identify cancer subtypes, tissues of origin, or predict survival^9–11^. Additional labels in the field of cancer biology may include the presence or absence of cancer, or response to therapy, and sparse genomic measures may be somatic mutations, circulating DNA fragments, neopeptides, RNA/protein modifications, copy number alterations, or methylation sites.

Somatic mutations are a complex, but well-studied genomic measure, with much of the biology already understood and ample data to test new models. When constructing features for somatic mutations our current understanding of the biology can easily be brought in, such as utilising information about genes^12–14^, or pathways^5^. However, for a given task it may not always be clear what known biology applies, and some measures may have uncertain biology. In these cases we can utilise the fundamental properties of the measure, and allow the model to show us which features are important through attention to specific instances and/or the learned representations of the instances. Some fundamental properties of a somatic mutation mutation are its local sequence context, which has been previously summarised by looking at the neighbouring 5’ and 3’ nucleotides^15,16^, and its genomic location, which has been represented as 1Mb bins^17^.

Here we present a tool for performing attention MIL, and demonstrate its application to somatic mutation data. We use this model to calculate attention for the fundamental properties of mutations, either local sequence context or the genomic position. Using simulated data we explore various MIL implementations on a range of tasks, and compare the proposed approach to conventional machine learning approaches in this area. We then apply the model to tumour classification, and learn the salient features of sequence and position while exceeding the performance of the current approaches. Finally we compare our model to state-of-the-art techniques at determining microsatellite status, and our model performs favourably despite the fact that comparable tools utilise a priori knowledge specific to the task while the proposed approach does not.

## RESULTS

### Aggregation tool for genomic concepts

Current applications of machine learning to mutation data are generally limited to an aggregation of hand-crafted features. Our model differs in that attention is first given to individual instances prior to aggregation, and if desired an end-to-end model is possible allowing for instance features to be learned during training (known as representation learning). Hand-crafted feature engineering may be most efficient when the representation of an instance is well/completely understood and aggregation of information over a set of instances is done via a static function such as sum and mean. Representation learning at the level of the instance can extract features specific to a given machine learning task, and in scenarios with a very large possible set of genomic measures can be combined with trainable attention mechanisms that can help with model explainability. To allow a model to extract its own features decisions must be made on how the raw data will be presented to the model, and how the model will encode the data. We consider the outcome of this process to be a genomic concept, and essential to extracting relevant features. Somatic mutations are often reported in a Variant Call Format (VCF) or Mutation Annotation Format (MAF), and a genomic concept can be constructed for any measurement in these files. The concept can be as simple as an embedding matrix (for example our position encoder), or it can be as complex as convolutional layers for the flanking nucleotide sequences along with the reference and alteration in both the forward and reverse directions (our sequence encoder, Figure 1).

To confirm that the encoders we developed were valid we performed positive controls, utilising the unique mutation calls from the TCGA MC3 public MAF. Our sequence encoder was found to be a faithful representation of a variant, learning the 96 contexts and an outgroup with near perfect accuracy, and our embedding strategy was able to perform a data compression without any information loss (Supplemental Figure 1). To confirm that our sequence encoder could effectively utilise strand information we asked a more difficult question, whether it could classify variants according to their consequence as provided by the MC3 MAF, specifically the consequence/variant classification of: frameshift deletion, frameshift insertion, inframe deletion, inframe insertion, missense, nonsense, silent, splice site, noncoding (5’ UTR, 3’ UTR, intron). This problem requires learning all 64 codons in 6 different reading frames, and importantly the strand the variant falls on affects the label. We first asked how well the model could do without providing reading frame, and while the model was able to learn InDels and splice sites, it was unable to distinguish noncoding mutations from the other classes (as would be expected), and did the best it could at associating codons with a consequence (Figure 2A). When provided the reading frame in the form of strand and CDS position modulo 3 (noncoding variants were represented by a zero vector), the sequence concept was now able to correctly classify missense vs. nonsense vs. silent mutations (Figure 2B), indicating that the modelling approach is able to learn which strand a feature was on, and correctly associate the relevant codons with consequence.

**Figure 2.**
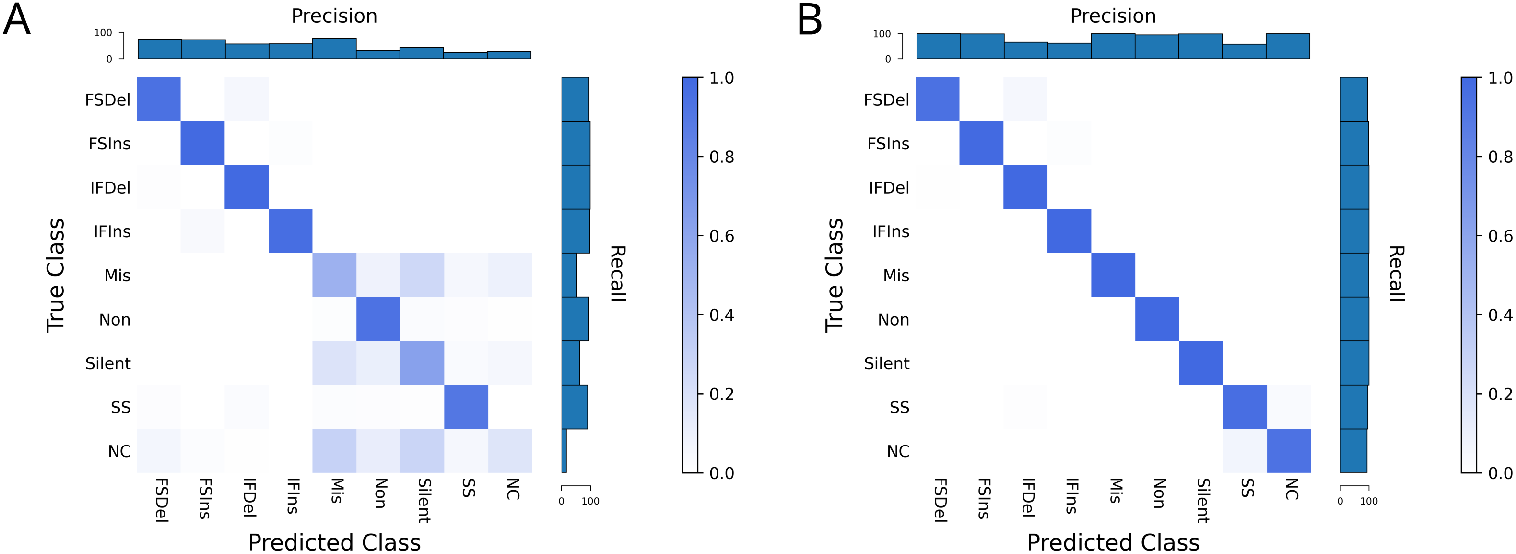
Predicting variant consequence. Our sequence encoder can learn variant consequence as defined by variant effect predictor (VEP) without (A) and clearly better with reading frame information (B). *FSDel=Frameshift deletion, FSIns=Frameshift insertion, IFDel=InFrame Deletion, IFIns=InFrame Insertion, Mis=Missense, Non=Nonsense, SS=Splice Site, NC=noncoding*. All four plots show row normalized confusion matrices.

Our implementation of MIL is motivated by Ilse et al.^18^ (Supplemental Figure 2), with some important modifications for the nature of somatic mutation data. In image analysis the aggregation function is often a weighted average, but whereas the number of tiles is unrelated to the label for an image the number of mutations may provide information about a tumour sample. Using simulated data we explored various MIL implementations along with traditional machine learning approaches, and found our attention-based MIL with a weighted sum performed well (Supplemental Figures 3-5). In order for a weighted sum to be meaningful the instance features must be activated, and this results in potentially large values on the graph. To account for this we perform a log of the aggregation on graph. We also developed dropout for MIL, wherein a random subset of instances is given to the model each gradient update, but then all instances are used during evaluation. This can allow for training with large samples, and also helps with overfitting since the samples are altered every batch. To improve model explainability we designed the model for multi-headed attention, where each attention head can be viewed as class-specific attention when the number of attention heads matches the number of classes.

### Cancer type classification

A readily available label in cancer datasets is the cancer type, and this task can have practical importance for when the tumour of origin for a metastatic cancer cannot be determined^19^. The types of mutations of a cancer are influenced by the mutational processes of its aetiology while the genomic distribution of its mutations is influenced by histology of origin, and these features of somatic mutations have been shown to be capable of classifying cancers^17,20–22^. We were interested in seeing how a model that calculates attention or learns it own features compared to established approaches. For this task we used the TCGA MC3 public mutation calls, which are exome-based. To understand the baseline performance on this data using current approaches we used two common hand-crafted features: the 96 SBS contexts (and a 97th outgroup so that no data was discarded), and an approximately 1Mb binning of the genome. For each manual feature we ran a logistic regression, random forest, neural net, and our MIL model. We also ran our model with 6 base pair windows of the sequences and explored gene as an input.

Table 1 shows the results of test folds from five-fold cross validation for the different models and the different inputs for the TCGA project codes (a 24 class problem). For each input our model outperformed current approaches, and the novel input of base pair windows showed a significant benefit (*∼*14% increase over the best standard model). Notably, our model showed a benefit even when using an identical encoding to the other models (96 contexts), suggesting the attention alone can improve model performance to some degree. To validate these results we investigated the performance of the 96 contexts in whole genome sequencing and again saw a benefit with our model (86% accuracy compared to 81%, Supplemental Table 1). For additional validation we also ran the models classifying the MC3 samples according to their NCIt codes and again saw a benefit with our model for every input (a 27 class problem, see Supplemental Table 2).

**Table 1.**
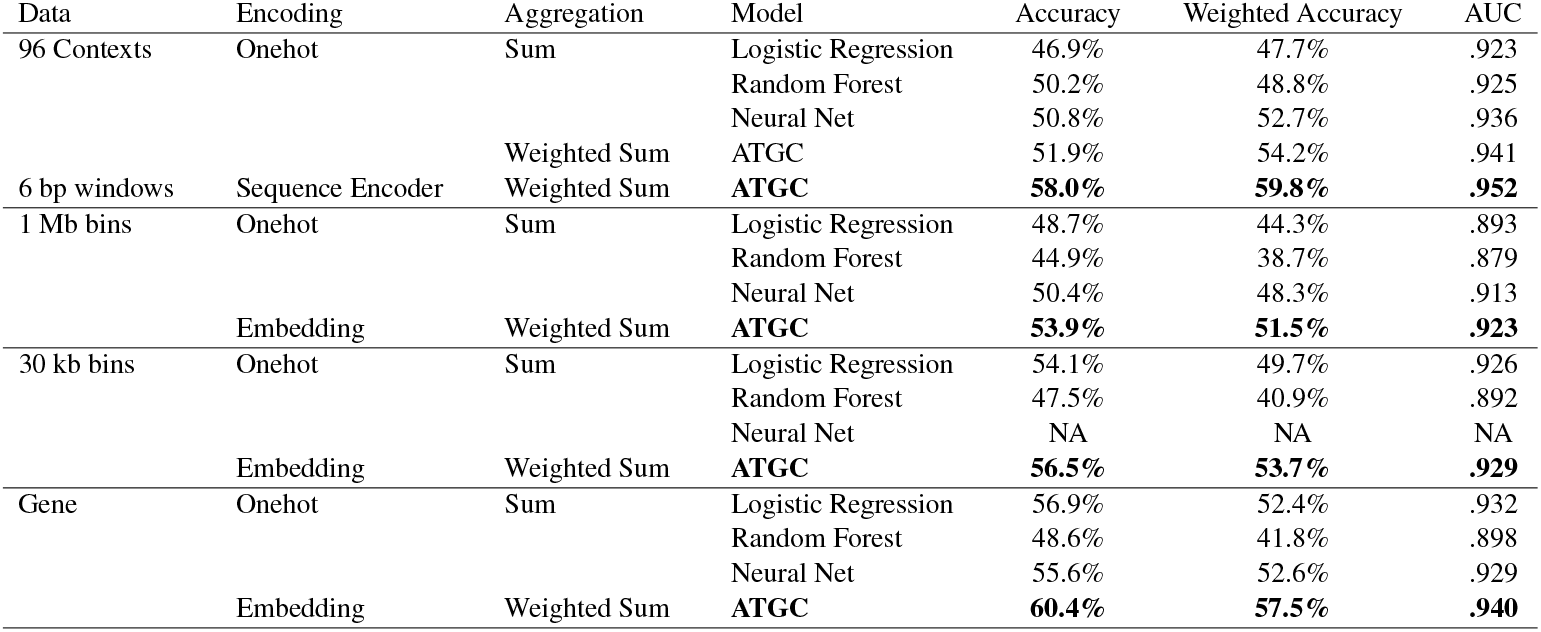
Tumour classification performance metrics for exome project codes. Every model was trained with the same sample weighting and 5-fold cross validation. When using a large input vector like the 30 kb bins our procedure for optimising neural nets cannot be used.

To investigate the performance differences we looked at classification performance in terms of precision and recall for each model stratified by tumour type (Figure 3A). Our model which takes 6 bp windows as input showed significant improvement in esophageal carcinoma (ESCA), lower grade glioma (LGG), pancreatic adenocarcinoma (PAAD), and testicular germ cell tumors (TGCT). The improvement seen in LGG is almost certainly due to identifying IDH1 mutations via a mapping of local sequence context of that specific hotspot. We also investigated the predictions of the best performing model (ATGC with 6 bp windows) with a confusion matrix (Figure 3B), and observed that cancers of similar histologic origin were often mistaken for each other. For the corresponding analyses with gene as input see Supplemental Figure 6.

**Figure 3.**
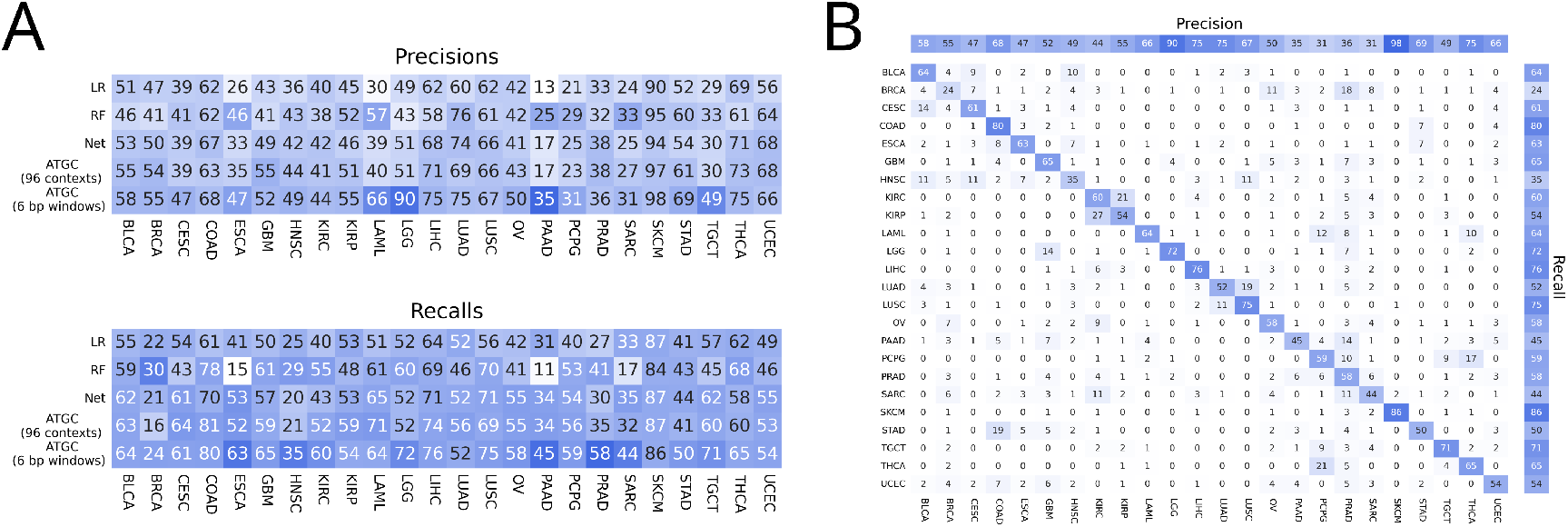
Tumour classification metrics. (A) Precisions and recalls for the models using the 96 contexts and ATGC with the 6 bp windows. (B) Confusion matrix for ATGC and the 6 bp windows.

When investigating how the model is interpreting genomic variant data we can examine both the representation of a variant produced by the model and what degree of attention the model is assigning a variant. To illustrate these two concepts we show a heatmap of the learned variant representation vectors for several cancer types with known aetiologies (Figure 4). Unsupervised K-means clustering was used to group the instances within each tumour type, revealing a rich feature space for the learned variant representations and clear clusters for each cancer type. We next explored how class specific attention related to this instance feature space and observed a spectrum of attention levels. In Figure 4 we see that for SKCM 6 clusters did for the most part produce clusters which were either high or low in attention, while in LUAD and COAD we see clusters that contain a bimodal distribution of attention.

**Figure 4.**
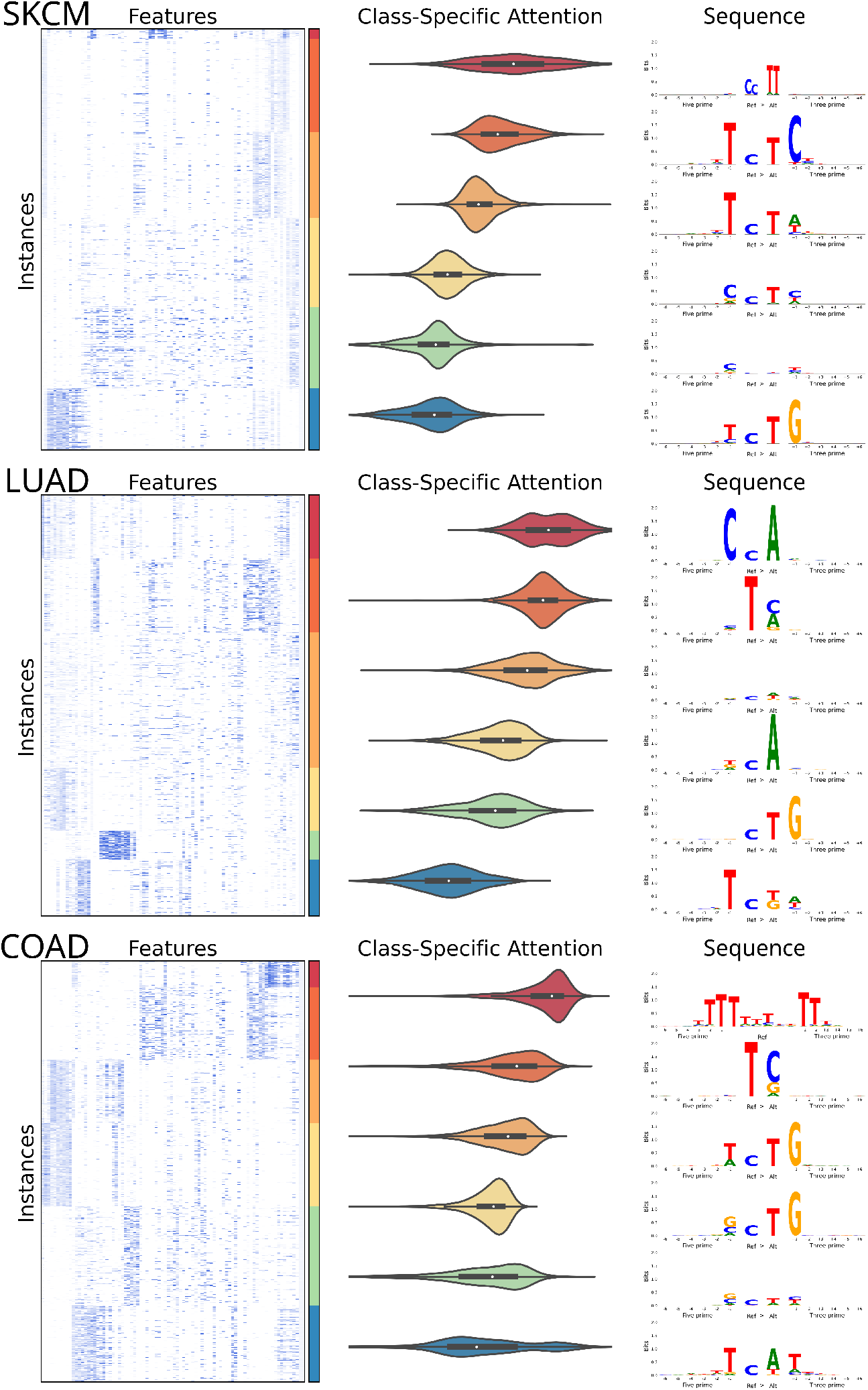
Instance feature vectors reveal known cancer biology. The feature vectors for the instances of a single test fold for SKCM, LUAD, and COAD samples were clustered with K-Means into 6 clusters and ordered by their median class specific attention (2000 instances sampled at random per cancer shown in the heatmaps). Violin plots of the attention values for each cluster are also shown along with sequence logos of the instances that make up each cluster.

To investigate what sequences were present in each of these clusters we generated sequence logos. For SKCM the highest attention cluster was composed of a specific doublet base substitution characteristic of ultraviolet radiation, with the next highest attention cluster comprised of a very specific 5’ nucleotide, ref, alt, and 3’ nucleotide also characteristic of ultraviolet radiation^16^. For LUAD the highest attention cluster contained sequences characteristic of tobacco smoking, while in COAD the highest attention cluster was a specific deletion occurring at a homopolymer, which is characteristic of cancers deficient in mismatch repair and is a known signature of this cancer type^16^.

While in Figure 4 we used clustering to identify groups of mutations within a cancer type which may be of interest, we instead could have simply sorted all instances by attention. To explore this possibility we took the highest attention instances (top 5%) of each attention head for SBSs, insertions, and deletions, and looked at the bits of information contained in the logos (Supplemental Figure 7). For SBS mutations most of the information gain occurs at the alteration and flanking nucleotides, however for most heads there is still information 2, 3, or more nucleotides away from the alteration. Given that a head of attention may identify several distinct motifs as important, the high attention instances for each head may not be homogeneous and as a result these bits represent a lower bound of the information gain. For InDels there is significant information 3 or 4 nucleotides away from the alteration, and these sequences are often mononucleotide repeats. When performing a similar analysis for our gene encoder we noticed that a small number of genes appear to be given high attention in each head, and clear groupings were present in the embedding matrix, with a small cluster enriched in cancer associated genes (Supplemental Figure 8).

Through the attention mechanism we also investigated what our model learned about genomic location. When looking at the attention values for the 1 Mb bins the different heads of attention appeared to be giving attention to the same bins, so we calculated the z-scores for each head and averaged across heads. We also noticed the values appeared consistent across folds of the data so we also averaged over the 5 data folds. Figure 5A shows the averaged attention values for the bins across the genome along with an embedding matrix from one of data folds. Specific bins are clearly either being upweighted or downweighted. Investigation of bins with low attention scores revealed that they contained very little data, which caused us to wonder if attention simply correlated with amount of data in each bin. There is initially a strong correlation between attention and number of mutations in a bin (Figure 5B), but once bins contain a certain amount of data the relationship disappears. Given that this is exomic data we suspected the bins with the highest attention contained genes important for cancer classification, so we calculated average gene attention z-scores with our model that used gene as input and matched genes with their corresponding bin (averaging when a gene was split across bins). There is nearly an order of magnitude more genes than bins, so most bins contain multiple genes and the same attention will be assigned to all genes in the bin. For the genes with the highest attention there is a corresponding bin also being given high attention (Figure 5C, top right corner), but there are also many genes which are passengers in those bins and mistakenly being given attention (Figure 5C, top left corner). This likely explains why using gene as input outperformed models using position bins as input.

**Figure 5.**
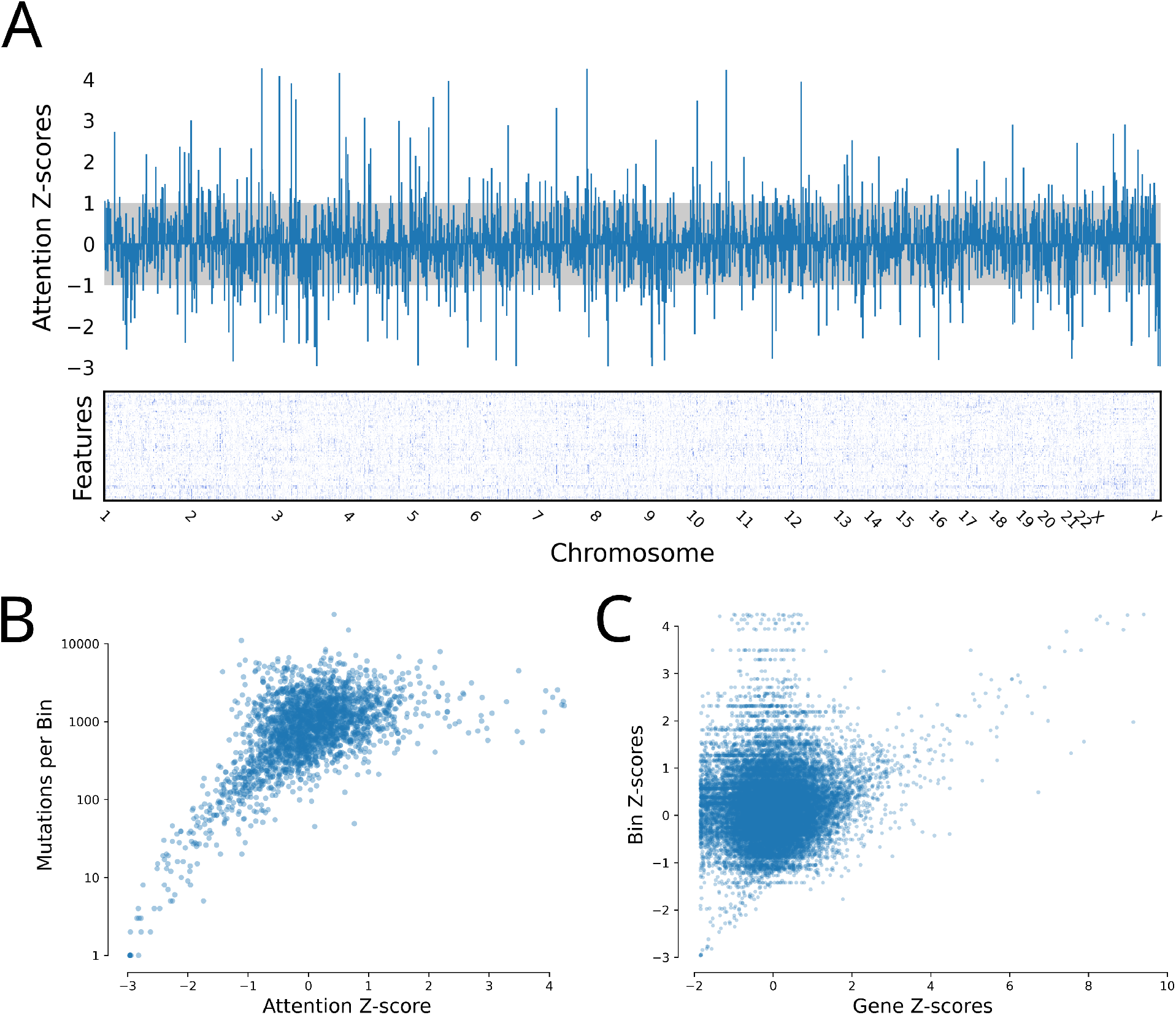
Genomic attention scores. (A) Attention z-scores for each 1 Mb bin were calculated across all 24 attention heads and averaged across 5 folds. The embedding matrix for a single fold is shown below with the starts of each chromosome shown. Z-scores between -1 and 1 are shaded. (B) Attention z-scores for each 1 Mb bin plotted against the number of mutations falling in each bin. (C) Attention z-scores for each 1 Mb bin plotted against the attention z-scores for the corresponding genes in those bins from a model with gene as input.

#### Microsatellite instability

Characterised by deficiencies in the mismatch repair proteins (MLH1, MSH2, MSH6, PMS2), microsatellite unstable tumours accumulate InDels at microsatellites due to polymerase splippage. As with tumor classification, current bioinformatic approaches to predicting MSI status rely on manually featurising variants. For example, the MANTIS tool uses a BAM file to calculate the average difference between lengths of the reference and alternative alleles at predefined loci known a priori to be important^23^. Similarly the MSIpred tool calculates a 22 feature vector using information from a MAF file and loci again known to be important, specifically simple repeat sequences^24^. Both of these approaches are valid and produce accurate predictions, but they rely on knowing the nature of the problem.

Because mutations characteristic of MSI occur at many different repeat sites, and because the repeats have a distinctive sequence, we chose to use our sequence concept for this problem. For the data we opted to use the controlled TCGA MC3 MAF rather than the public MAF since the public MAF excludes most variants in non-exonic regions and most simple repeats fall in these regions. The TCGA has ground truth labels as defined by PCR assay for some tumour types, and we were able to obtain labels for uterine corpus endometrial carcinoma (UCEC, 494), stomach adenocarcinoma (STAD, 437), colon adenocarcinoma (COAD, 365), rectum adenocarcinoma (READ, 126), ESCA (87), and uterine carcinosarcoma (UCS, 56) tumour samples. For the sequence concept we went out to 20 nucleotides for each component (5’, 3’, ref, and alt) to allow the model to potentially capture long repetitive sequences.

Although we did not provide the model information about cancer type or perform any sample weighting, our model displayed similar performance across cancer types (Supplemental Figure 9A). We believe MANTIS and MSIpred are considered state of the art when it comes to MSI classification performance, and as can be seen in Figure 6A our model slightly performs them despite not being given information about simple repeats. When comparing our predictions to MANTIS both models were often similarly confident in their predictions (Supplemental Figure 9B), however there are cases where MANTIS predicts the correct label but our model doesn’t, or our model predicts the correct label but MANTIS does not, suggesting that the best MSI predictor may be one that incorporates predictions from multiple models. There are a few cases where the sample is labelled MSI high by the PCR, but both MANTIS and our model are confident the sample is MSI low, perhaps indicating the PCR label may not be 100% specific (or alternatively indicates an issue with the BAM files for these samples).

**Figure 6.**
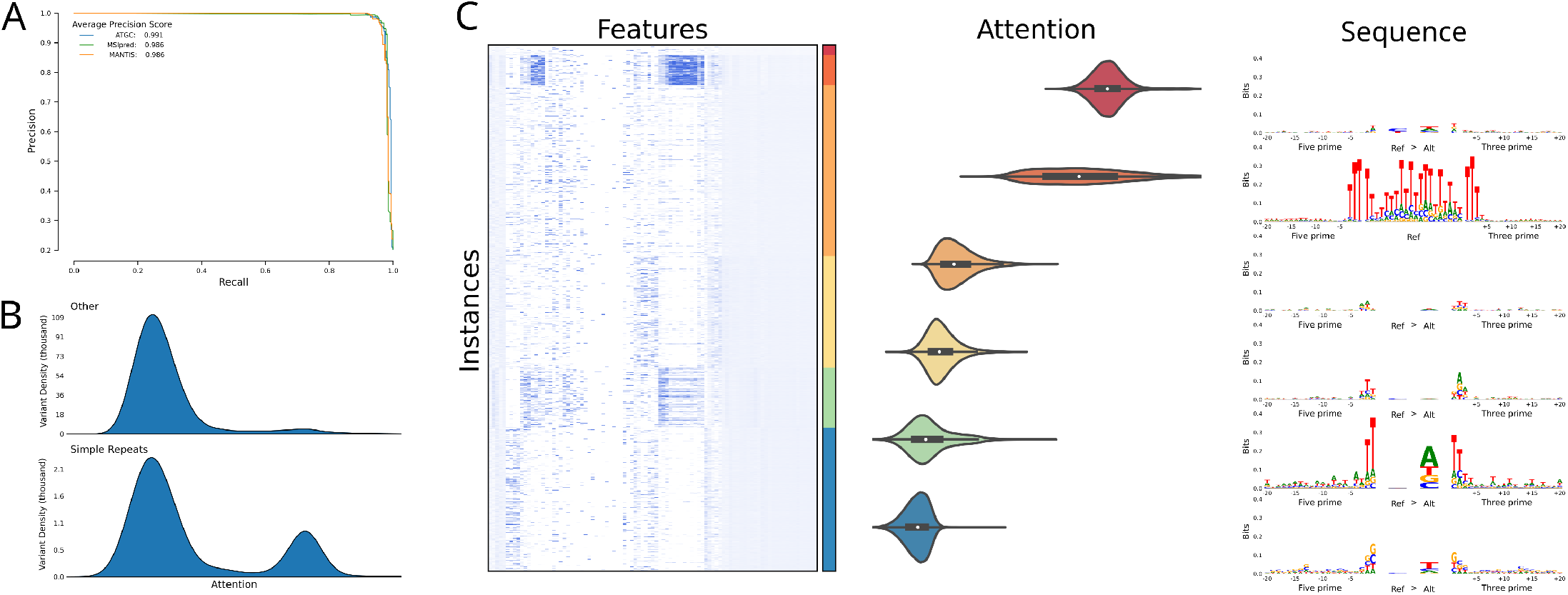
Application of ATGC MIL framework accurately predicts MSI status by learning which are the salient variants. (A) Average precision recall curves over 9 stratified K-Fold splits for MSIpred and ATGC, precision recall curve for MANTIS for samples which had a MANTIS score. (B) Probability density plots of the output of the attention layer stratified by whether a variant was considered to be in a simple repeat from the UCSC genome annotations. (C) Features of 2000 random instances from a single test fold clustered with K-Means and the corresponding sequence logos of the clusters.

To classify whether a variant falls in a repeat region or not MSIpred relies on a table generated by UCSC (simpleRe-peat.txt). Using this file we labelled variants according to whether they fell in these regions, and as seen in Figure 6B variants which occurred in a simple repeat region were much more likely to receive high attention than variants that did not. When clustering the instances of these samples we observed two clear clusters which are receiving high attention, a smaller cluster characterised by SBS mutations in almost exclusively intergenic regions with almost 33% labelled as a simple repeat, and a larger cluster characterised by deletions in genic but noncoding regions. The sequence logo of the intergenic cluster did not reveal a specific sequence while the logo for the deletion cluster revealed that the model is giving attention to deletions at a mononucleotide T repeat.

## DISCUSSION

Many genomic technologies generate data which can be considered “large p, small n”, wherein the number of possible measures/features per sample greatly exceeds the number of samples. For example, somatic mutations can occur anywhere in the genome, thus creating an enumerable number of possible unique features per sample. Similar considerations apply to circulating DNA fragments, CHIP-SEQ peaks, methylation sites, or RNA/protein modifications. Attention MIL is a natural solution to these problems because it essentially transposes the problem—the large amount of instance data is a benefit instead of a hindrance when extracting relevant features.

When performing cancer classification, application of our model led to improvement regardless of the input; and when classifying samples according to their MSI status our model slightly outperformed the current state-of-the-art methods despite the other methods containing information specific to the task. Importantly, whereas other models require post hoc analyses to understand how they are making decisions, our tool directly reveals which instances it views as important. This is essential because as these measurements begin to be used in the clinic and researchers turn to deep learning for their analysis, it will likely be necessary for the models to explain their decisions given a patient’s right to understanding treatment decisions^25^.

Research into how to best implement attention MIL is an active field, with a recent activity in its application to computational pathology^26^. We consider our approach to attention MIL fairly standard^18^, with our goal being to simply demonstrate its value in the context of genomics data. As such, it is unlikely our results represent the full potential of applying MIL to this data. For example, in computational pathology there have been recent suggestions to include a clustering step at the instance level^9,27,28^. And by design we limited our instance featurisation to the factors that uniquely define a mutation: chromosome, position, and reference/alternative alleles. We can easily imagine incorporating outside knowledge into the variant encoding process, such as variant consequence, biological pathway(s) involved, etc. We hope that the success we achieved with our proof-of-concept application of MIL inspires additional work in this area.

## METHODS

### Model

We used a combination of TensorFlow 2 and tf.keras for implementing ATGC. Keras, like many deep learning libraries, requires the first dimension of the inputs to match the first dimension of the outputs. Many implementations of MIL work around this constraint by performing stochastic gradient descent, where one sample is shown to the model at a time. This precludes the ability to easily perform sample weighting. To perform minibatch gradient descent as we would with any other model we developed our model around ragged tensors.

Our model is modular in the sense that the top aggregation model which calculates attention and generates predictions takes as input encoder models. These encoders perform operations on ragged data for instance data and normal vectors for sample data (the dataset functions automatically infer whether a ragged tensor needs to be made). This framework allows for any number of instance or sample inputs, any number of outputs, and any number of attention heads (as GPU memory permits). Ragged tensors are fully supported by TensorFlow, so it is possible to use our model with default loss functions and dataset batching. However, because we like additional control over the sample weighting and want the ability to perform stratified batching and data augmentation (data dropout), we prefer to use our own loss and metric classes even when the loss and/or metric already exists in TensorFlow, and we created dataset utilities built around generators.

Our attention is inspired by the attention first proposed by Ilse et al.^18^. One issue with the proposed attention in MIL is that its value is not only a function of an instance’s importance, but also its rarity. If two instances are equally important, but one is present at a much lower fraction (low witness rate), then the rarer instance will be given a higher attention, and if a key instance is very frequent it may be given very little attention if any. In addition it’s possible for the model to find a solution that is characterised by the absence rather than presence of key instances, which results in key instances being given a lower attention. To correct for this issue and make the attention clearer, we added L1 regularisation on the output of the attention layer. The more regularisation added the clearer the separation between key instances and negative instances, but at the risk of decreased model performance. Another potential issue with attention is that it is independent of the bag. It may be the case that a key instance is only a key instance when it occurs in a certain bag environment, but the model will not make this distinction at the level of the attention layer. If desired, the attention can be made dynamic^29^ by sending information about the samples back to the instances (Supplemental Figure 2C), which may provide additional information about how the model is making decisions. We developed our own version of dynamic attention which calculates a weighted mean with standard MIL attention, then sends that sample vector back to the instances and calculates a second round of attention and performs a second aggregation.

We provide users several options for the aggregation function (mean, sum, dynamic). When using an aggregation function that includes a sum the instance features should be activated, and this can result in potentially large sums. To counteract this we log the aggregations on the graph when a sum is performed. Depending on the data there may be too many mutations to fit onto the graph, so we developed dropout for attention MIL where a random fraction of instances per batch are sent into the model during training, but then all the instances are used for evaluation. This has the additional benefit of helping with overfitting as the samples are constantly changing during training, and is essentially a form of data augmentation.

### Custom activation functions

Functions can be tested at: https://www.desmos.com/calculator/vruk2tomxk

#### Adaptive square root (ASR)

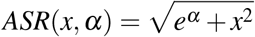

The above will be used as a core element in the activation functions below.

#### Adaptive rectifying unit (ARU)

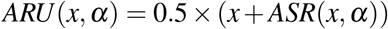

Note, a bias term could be added and applied to *x* prior to this activation function (as is the convention) and *α* can be a trainable parameter that modulates the curvature of this “rectifier”. The ARU can approach the shape of the commonly used rectified linear unit (ReLu) as *α* approaches negative infinity, but ARU is fully differentiable across all *x*, which does not hold for ReLu. We have tuned default initial conditions for *α* to match the commonly used softplus function, particularly near *x* = 0, as initial conditions for this adaptive rectifier.

#### Adaptive sigmoid unit (ASU)

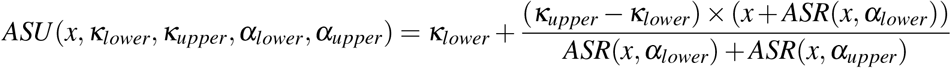

Again, a bias term could be added and applied to *x* prior to this activation function (as is the convention). We generally keep the lower and upper asymptotes fixed at 0, 1 or –1, 1 depending on the application. The lower and upper *α* parameters together control the curvature of the sigmoid. We have tuned default values to match the logistic function, particularly near *x* = 0, as initial conditions for this adaptive sigmoid function.

The above were motivated by the previously described inverse square root unit (ISRU)^30^. It is important to note that all input *x* is at most squared in this formulation and there is no application of input *x* as an exponent (*e*^*x*^), which greatly helps to avoid numerical overflow that can occur in the context of aggregation over samples in a multiple instance learning (MIL) framework.

### MAF processing

For the MC3 public MAF variants which had a “FILTER” value of either “PASS”, “wga”, or “native wga mix” and fell within the coordinates of the corresponding coverage WIGs were retained. For the MC3 controlled MAF variants which had a “FILTER” value of “PASS”, “NonExonic”, “wga”, “bitgt”, “broad PoN v2”, “native wga mix” or combination thereof and were called by more than one mutation caller were retained. The MC3 working group was inconsistent in its merging of consecutive SBSs, so these were merged into a single mutation if the maximum difference between the average alternative or reference counts and any single alternative or reference count was less than 5 or the maximum percent difference was less than 5%, or the maximum VAF deviation was less than 5%.

### Tumour classification analyses

We performed tumour classification with both the original project codes and an NCIt ontology for exomic data, and with histology codes for whole genome data. For the project codes we required each class to have 125 samples, resulting in 24 classes and 10,012 samples. For the NCIt classification, tumours were mapped to their NCIt code by Alex Baras, a board certified pathologist, using the available histology description. We required at least 100 samples per NCIt code, resulting in 27 classes and 8,910 samples. For the PCAWG data only white-listed samples were used and a donor was only allowed to have a single sample in the dataset. When a donor had more than one sample the following preference order of specimen type was used for selection of the sample: Primary tumour - solid tissue, Primary tumour - other, Primary tumour - lymph node, Primary tumour - blood derived (peripheral blood), Metastatic tumour - metastasis local to lymph node, Metastatic tumour - lymph node, Metastatic tumour - metastasis to distant location, Primary tumour - blood derived (bone marrow). To prevent potential issues with GPU memory we only used samples with less than 200k mutations. Any histology with at least 33 samples was used for analysis, resulting in 24 histologies and 2,374 samples. All models were weighted by tumour type, and the K-Folds were stratified by tumour type.

#### Logistic regressions

“LogisticRegression” from “sklearn.linear model” was used with default parameters.

#### Random forests

We used “RandomForestClassifier” from “sklearn.ensemble” and we explored how the different parameters affected performance with “gp minimize” from “scikit-optimize”. We found most parameters to have limited effect, however we did find the default number of estimators was far from optimal, and set “n estimators” to 900 and “min samples split” to 10.

#### Neural nets

The process from the PCAWG consortium was used for constructing the neural nets^17^. Essentially for each fold of the data 200 different sets of hyperparameters are searched through. Since we perform weighted training we selected the best performing hyperparameters based on weighted cross entropy instead of accuracy. When using gene as input the default search space caused some issues with GPU memory and as a result were adjusted, but otherwise the search space was copied exactly.

#### MIL models

Details for each individual MIL model that was run can be found in the accompanying GitHub repository.

### MSI analyses

The PCR labels were merged and when 5 marker call and 7 marker call were in disagreement, the 7 marker call was given preference. Both MSS and MSI-L were considered to be MSI low. The source code for MSIpred was altered to allow it to run on Python 3, and the model was changed to output probabilities, but otherwise was run as recommended. Models were not weighted by tumour type, but tumour type was used for K-Fold stratification. For our MIL model because there were only 2 classes we considered it a binary classification task and used a single head of attention. A data dropout of .4 was used, and the sequence components were given 8 independent kernels with ARU activation and fused to a dimension of 128 with a relu activation and .01 L2 kernel regularisation, a dropout of .5 was performed, then attention was calculated with a single layer and ASU activation and .05 L1 activity regularisation to force the positive class to receive higher attention. Following aggregation a layer of 256 and relu activation was used followed by .5 dropout, a layer of 128 and relu activation followed by .5 dropout, to the final prediction and no activation.

### Sequence logos

Logomaker^31^ was used for the sequence logos. The probabilities were calculated separately for single base substitutions, doublet base substitutions, insertions, and deletions.

## DATA AVAILABILITY

All data used in this publication is publicly available. The MC3 MAFs are from Ellrott et al.^32^, the MSI PCR labels are available from cBioPortal, TCGAbiolinks, or individual publications^33–36^. The UCSC simpleRepeat.txt was downloaded from http://hgdownload.cse.ucsc.edu/goldenPath/hg19/database/simpleRepeat.txt.gz. The Broad coverage WIGs are available at https://www.synapse.org/#!Synapse:syn21785741. MANTIS values are from Bonneville et al.^37^. The MAF for the ICGC PCAWG samples was obtained from https://dcc.icgc.org/releases/PCAWG/consensus_snv_indel, and the MAF for the TCGA PCAWG samples was obtained at https://icgc.bionimbus.org/files/0e8a845d-a4f4-40bc-890b-5472702d087c.

## CODE AVAILABILITY

All code for processing data, running models, and generating figures is available at GitHub: https://github.com/OmnesRes/ATGC/tree/method_paper, and has been archived at Zenodo: https://doi.org/10.5281/zenodo.8083498. Code is written in Python 3 and TensorFlow 2. All intersections were performed with PyRanges^38^. We leveraged NVIDIA V100s with 32GB of RAM for much of the computation, however most of the computations here could be reasonably performed on CPU as well within the same coding framework.

## ACKNOWLEDGEMENTS

The results here are in whole or part based upon data generated by the TCGA Research Network. J.A. and A.S.B. disclose support for the publication of this study from Mark Foundation for Cancer Research (19-035-ASP). A.S.B. discloses support from The Leon Troper, M.D. Professorship in Computational Pathology at Johns Hopkins.

## CONTRIBUTIONS

J.A., J.-W.S., and A.S.B. conceived the study. J.A. and A.S.B designed experiments. J.A. performed experiments and analysed results. F.M., and A.S.B supervised the research. J.A. and A.S.B. wrote the manuscript with all authors providing input.

## REFERENCES

1. Ching, T. et al. Opportunities and obstacles for deep learning in biology and medicine. Journal of The Royal Society Interface 15, 20170387 (2018).

2. Zhou, J. & Troyanskaya, O. G. Predicting effects of noncoding variants with deep learning–based sequence model. Nature Methods 12, 931–934 (Aug. 2015).

3. Routhier, E. & Mozziconacci, J. Genomics enters the deep learning era. PeerJ 10, e13613 (2022).

4. Altman, N. S. & Krzywinski, M. The curse (s) of dimensionality. Nature Methods 15, 399–400 (2018).

5. Elmarakeby, H. A. et al. Biologically informed deep neural network for prostate cancer discovery. Nature 598, 348–352 (2021).

6. Dietterich, T. G., Lathrop, R. H. & Lozano-Pérez, T. Solving the multiple instance problem with axis-parallel rectangles. Artificial Intelligence 89, 31–71 (1997).

7. Amores, J. Multiple instance classification: Review, taxonomy and comparative study. Artificial intelligence 201, 81–105 (2013).

8. Carbonneau, M.-A., Cheplygina, V., Granger, E. & Gagnon, G. Multiple instance learning: A survey of problem characteristics and applications. Pattern Recognition 77, 329–353 (2018).

9. Lu, M. Y. et al. Data-efficient and weakly supervised computational pathology on whole-slide images. Nature Biomedical Engineering 5, 555–570. https://doi.org/10.1038/s41551-020-00682-w (2021).

10. Lu, M. Y. et al. AI-based pathology predicts origins for cancers of unknown primary. Nature 594, 106–110. https://doi.org/10.1038/s41586-021-03512-4 (2021).

11. Chen, R. J. et al. Whole Slide Images are 2D Point Clouds: Context-Aware Survival Prediction Using Patch-Based Graph Convolutional Networks in Medical Image Computing and Computer Assisted Intervention – MICCAI 2021 339–349 (Springer International Publishing, 2021). https://doi.org/10.1007/978-3-030-87237-333.

12. Kim, S., Lee, H., Kim, K. & Kang, J. Mut2Vec: distributed representation of cancerous mutations. BMC Medical Genomics 11, 33 (2018).

13. Palazzo, M., Beauseroy, P. & Yankilevich, P. A pan-cancer somatic mutation embedding using autoencoders. BMC Bioinformatics 20, 655 (2019).

14. Peng, J., Zou, D., Gong, W., Kang, S. & Han, L. Deep neural network classification based on somatic mutations potentially predicts clinical benefit of immune checkpoint blockade in lung adenocarcinoma. OncoImmunology 9, 1734156 (2020).

15. Alexandrov, L. B. et al. Signatures of mutational processes in human cancer. Nature 500, 415–421 (2013).

16. Alexandrov, L. B. et al. The repertoire of mutational signatures in human cancer. Nature 578, 94–101 (2020).

17. Jiao, W. et al. A deep learning system accurately classifies primary and metastatic cancers using passenger mutation patterns. Nature Communications 11, 728. ISSN: 2041-1723 (2020).

18. Ilse, M., Tomczak, J. M. & Welling, M. Attention-based deep multiple instance learning. arXiv preprint arXiv:1802.04712 (2018).

19. Pavlidis, N. & Pentheroudakis, G. Cancer of unknown primary site. The Lancet 379, 1428–1435 (2012).

20. Salvadores, M., Mas-Ponte, D. & Supek, F. Passenger mutations accurately classify human tumors. PLoS computational biology 15, e1006953 (2019).

21. Danyi, A., Jager, M. & de Ridder, J. Cancer type classification in liquid biopsies based on sparse mutational profiles enabled through data augmentation and integration. Life 12, 1 (2021).

22. Sanjaya, P., Waszak, S. M., Stegle, O., Korbel, J. O. & Pitkanen, E. Mutation-Attention (MuAt): deep representation learning of somatic mutations for tumour typing and subtyping. bioRxiv (2022).

23. Kautto, E. A. et al. Performance evaluation for rapid detection of pan-cancer microsatellite instability with MANTIS. Oncotarget 8, 7452 (2017).

24. Wang, C. & Liang, C. MSIpred: a python package for tumor microsatellite instability classification from tumor mutation annotation data using a support vector machine. Scientific Reports 8, 1–10 (2018).

25. Goodman, B. & Flaxman, S. European Union regulations on algorithmic decision-making and a “right to explanation”. AI magazine 38, 50–57 (2017).

26. Gadermayr, M. & Tschuchnig, M. Multiple Instance Learning for Digital Pathology: A Review on the State-of-the-Art, Limitations & Future Potential. arXiv preprint arXiv:2206.04425 (2022).

27. Li, J. et al. A multi-resolution model for histopathology image classification and localization with multiple instance learning. Computers in biology and medicine 131, 104253 (2021).

28. Sharma, Y. et al. Cluster-to-conquer: A framework for end-to-end multi-instance learning for whole slide image classification in Medical Imaging with Deep Learning (2021), 682–698.

29. Yan, Y. et al. Deep multi-instance learning with dynamic pooling in Asian Conference on Machine Learning (2018), 662–677.

30. Carlile, B., Delamarter, G., Kinney, P., Marti, A. & Whitney, B. Improving deep learning by inverse square root linear units (ISRLUs). arXiv preprint arXiv:1710.09967 (2017).

31. Tareen, A. & Kinney, J. B. Logomaker: beautiful sequence logos in Python. Bioinformatics 36, 2272–2274 (2020).

32. Ellrott, K. et al. Scalable open science approach for mutation calling of tumor exomes using multiple genomic pipelines. Cell Systems 6, 271–281 (2018).

33. Cancer Genome Atlas Network et al. Comprehensive molecular characterization of human colon and rectal cancer. Nature 487, 330 (2012).

34. Levine, D. A. Integrated genomic characterization of endometrial carcinoma. Nature 497, 67–73 (2013).

35. Berger, A. C. et al. A comprehensive pan-cancer molecular study of gynecologic and breast cancers. Cancer cell 33, 690–705 (2018).

36. Liu, Y. et al. Comparative molecular analysis of gastrointestinal adenocarcinomas. Cancer cell 33, 721–735 (2018).

37. Bonneville, R. et al. Landscape of Microsatellite Instability Across 39 Cancer Types. JCO Precision Oncology, 1–15 (Nov. 2017).

38. Stovner, E. B. & Sætrom, P. PyRanges: efficient comparison of genomic intervals in Python. Bioinformatics 36, 918–919 (2020).

