## Supplemental Information for "Aggregation Tool for Genomic Concepts (ATGC): A deep learning framework for somatic mutations and other sparse genomic measures"

### GENOMIC CONCEPTS

Any aspect of a variant can be featurised (surrounding sequence, reference, alteration, location, gene, consequence, allele frequency, etc.), and the choice of how that featurisation is performed is what we consider a genomic concept. In this manuscript we focused on featurising the alteration and surrounding sequence, the position, and the gene the mutation occurs in.

#### Sequence concept

Our concept of the sequence of a somatic alteration splits a variant into its 5' sequence, 3' sequence, reference sequence, and alternative sequence. Because of the symmetry of DNA and the fact that it is impossible to know which strand a mutation occurred on, there are two equivalent representations for any given variant, and we stack these two equivalent representations on top of each other and allow the model to decide which representation to use. If the strand the variant is on has any relevance the model will use that information to decide which representation to use. While the sequences can be as long as desired, because large InDels must be clipped and we perform the clipping by taking the sequences at the edges, there is a requirement for those sequences to be an even length. It is currently thought that only neighbouring sequences contain any signal, and as a result we focused on shorter sequence lengths (up to 20 nt). Because the sequences are short, for computational efficiency we used a single convolutional layer to featurise each sequence component individually (5', 3' ref, alt), and then fused the resulting features for each representation separately. If a user is interested in longer sequences and specific motifs multiple convolutional layers can be used.

Variants when reported in a VCF contain empty alleles whereas there are no empty alleles in a MAF. As a result, if a model is trained on one format then it should only be applied to data of the same format. Similarly, when reporting an insertion or deletion in a repetitive region there are currently 2 formats for reporting these variants, either left or right aligned. With left alignment the repeat will be present in the 3' sequence, while with right alignment the repeat will be present in the 5' sequence. A model trained on left-aligned data would not be applicable to data that is right-aligned. To avoid this issue and to allow a repeat to be present in both the 5' region and 3' region we created a new alignment for variants, center-alignment (technically left-center alignment since a mutation can't be placed in the center of an odd number of repeats).

#### Position and Gene Concepts

Previously position in the genome has been represented by a genomic bin. This binning is essentially a categorical input similar to genes. In both cases the number of bins and genes is large, causing issues for neural networks and making it difficult to calculate attention in an MIL framework. We solve this problem by encoding these inputs with a trainable embedding. This compresses what would have been a large onehotted feature space to a user-defined space (for example 128). The embedding matrix also provides a level of interpretability—inputs with a similar embedding would be expected to have a similar contribution to the learning task.

### VALIDATION OF GENOMIC CONCEPTS

To confirm that our concepts are accurately encoding the relevant features of variants we first performed some positive controls. For the data we used the public MAF provided by The Cancer Genome Atlas (TCGA) MC3 working group. We extracted the reference and alternative sequences directly from the MAF, padding SBSs and InDels and clipping InDels where necessary. 5' and 3' sequences were pulled from the human genome (GRCh37) using the provided start positions. For each sequence element in the concept we used six nucleotides.

When defining a SBS with the pyrimidine as the reference sequence there are only six types of SBSs (T>C, T>G, T>A, C>T, C>G, C>A), and when considering the neighbouring five prime and three prime nucleotides there are $6\times4\times4$ = 96 contexts. It is these 96 contexts which have been used to define mutational signatures. We labelled the variants in the MAF with these 96 contexts if the variant was a SBS, and gave a separate label for other variant types, resulting in a total of 97 classes. Supplemental Figure 1A shows the results of the 97 class classification problem. Our concept achieved near perfect accuracy, giving us confidence that our sequence concept is a valid representation of a variant's alteration and neighbouring sequences.

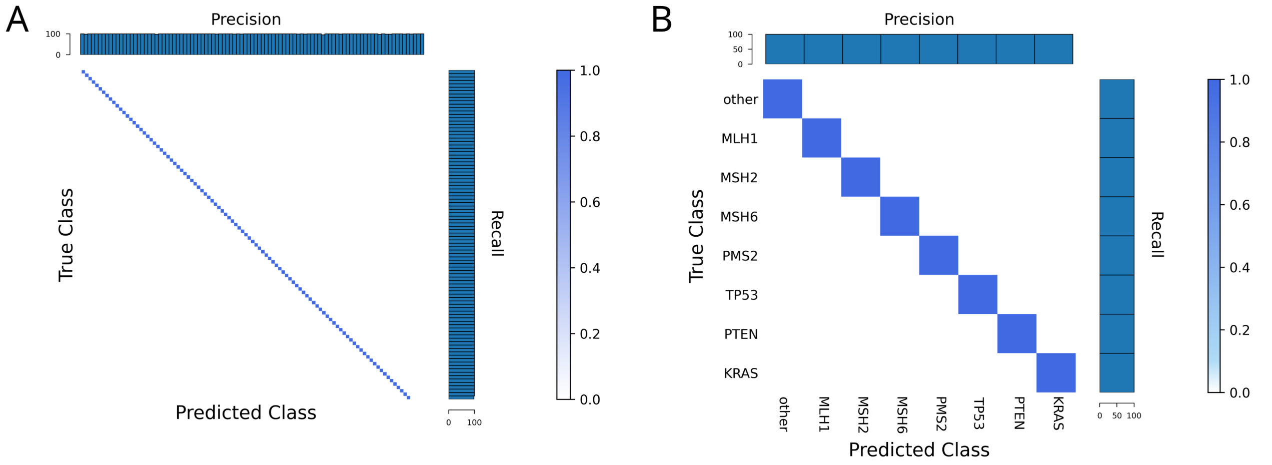


**Supplemental Figure 1. Characterisation of performance at the mutation (instance) level.** (A) 97 way classification (96 nucleotide contexts and an other/out group category) in which the model is able to accurately learn the conventional trinucleotide mutational contexts. (B) Our gene encoder can compress gene information with perfect accuracy.

For the gene concept we simply wanted to know whether it is possible to compress the feature space without any information loss. Supplemental Figure 1B shows that we were able to perfectly classify some genes of interest at the actual frequency they occur in the data with an embedding dimension of 128.

### SAMPLE-LEVEL SUPERVISED LEARNING: AGGREGATION OF GENOMIC CONCEPTS

In MIL, at some point the instance features for each bag must be aggregated into a single sample-level tensor that can then be used to compare against the bag label. There are two straightforward ways of accomplishing this: either bringing the instance features to the dimension of the output and then aggregating (the instance model, Supplemental Figure 2A), or aggregating first and then bringing the sample features to the output dimension (the sample model, Supplemental Figure 2B). The first approach coincides nicely with the traditional formulation of MIL—each positive bag is thought to contain one or more key instances and the negative bags do not contain these instances. The problem is essentially reformulated as classifying the instances and then pooling these classifications. The appeal of this approach is that the model makes it very clear which instances were thought to be associated with each class. However, if the problem of MIL is reenvisioned as the bag label being determined by global properties of the bag instead of single instances, for example a positive bag being defined as containing multiple types of instances or amounts of instances, then only the sample model will be able to identify these properties and correctly classify the bags. As a result, the sample model is considered to be a more general solution to the multiple instance problem, but the model no longer makes it clear which instances were deemed key instances.

To get the best of both worlds an attention can be given to each instance (Supplemental Figure 2C). The attention is derived directly from the instance feature vectors, can be single or multi-headed, and is multiplied back against the instance feature vectors as a weighting factor. The larger the value, the more important the model views an instance. In addition to giving the sample model a level of explainability, it improves model performance by upweighting the important instances and downweighting the unimportant instances (noise reduction).

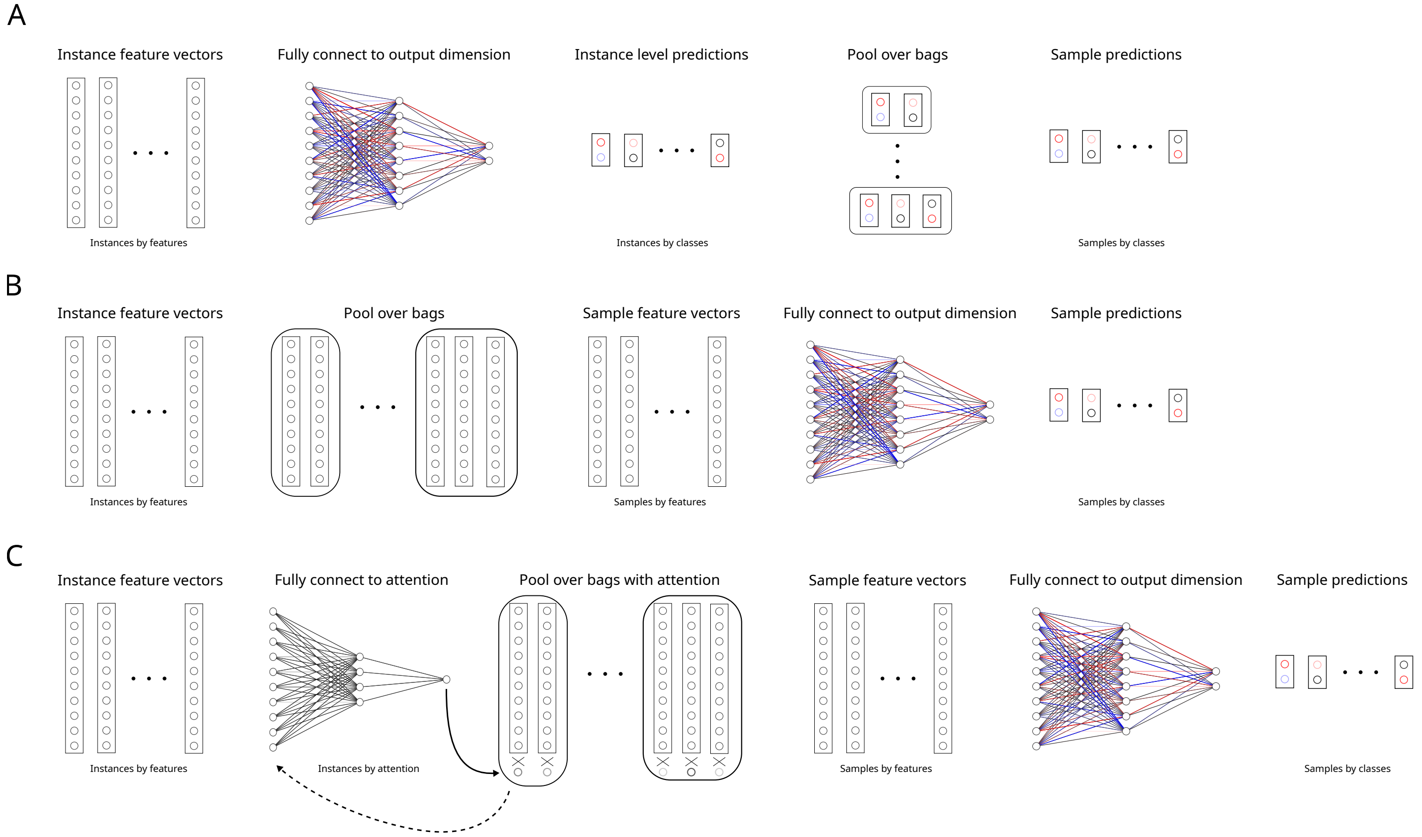


**Supplemental Figure 2. Multiple instance learning approaches.** (A) *MIL based on an instance classifier.* The instance feature vectors are used to determine the class of the instances, and then aggregated. (B) *MIL based on a sample classifier.* The instance features are first aggregated into a sample feature vector, and then the sample feature vectors are used to classify the samples. (C) *MIL with attention.* The instances are first reduced to a single dimension for each head of attention, each of which can be interpreted as the attention of an instance, and this attention is multiplied against each corresponding instance vector. The resulting weighted instance feature vectors are then aggregated, and the resulting sample features are then used to classify the samples. It is possible to send the sample features back to the instances and calculate the attention a second or more time(s).

### Performance Characteristics of Different Aggregation Methods

Witness rate, bag size, bag composition, number of training examples, problem type (regression vs. classification), label noise, unrepresentative negative space, etc., can all uniquely affect the performance of a MIL model. Since MIL has never been applied to sparse genomics data before we systematically explored various MIL approaches across a range of tasks to try and understand the performance characteristics of each paradigm. Regardless of the approach used, due to the order of the instances of each bag being irrelevant, the aggregation must be a permutation-invariant operation to summarise the contents of each bag. Some examples of typical operations used include mean, sum, and max, with max being particularly relevant when the witness rate is low. For the purposes of our simulated problems we focused on comparing mean and sum as they seemed most suited for the tasks.

In addition to taking a weighted mean or weighed sum where the weighting (attention) is independent of the bag, it is possible to have the attention be dependent on the bag. We refer to our bag-dependent attention as the "dynamic" model, whereby a weighted average is sent back to the instances to calculate a second attention which is then used to calculate a weighted sum. To compare against the current state-of-the-art machine learning methods for somatic variants we also applied random forests and logistic regression to aggregations of the 96 contexts.

For all simulated data the bag sizes are random, albeit an attempt was made to have the bag size distribution somewhat resemble what one might expect in panel data (with the important exception of each bag having at least one variant). In all cases 1000 samples were generated, with 500 used for training, 300 for validation, and 200 for test. For the classification experiments approximately half of the bags were positive, and half were negative. As a result, a random classifier would have 50% accuracy. Key instances were always variants with a specific 5' sequence, and negative variants were randomly generated (and checked to not have the 5' sequence of key instances). The encoder for each model was our sequence encoder. None of the experiments were meant to be particularly hard (witness rate was generally high and there was limited label noise added if any), but that fact underscores a model's inability to solve a problem when it doesn't perform well.

#### Classification Experiments

We performed four different classification experiments on all the models (Supplemental Figure 3). The first experiment corresponds to the classical formulation of MIL: the positive bags contain one or more key instances not observed in the negative bags. As a result, we expected all models which could identify key instances to succeed. The MIL models all solved the problem while the machine learning methods could not. The MIL models with attention all gave attention to the key instances.

In the second experiment we diverged from the classical MIL formulation, and required the model to learn a bag-level property: the presence of two different key instances. Because the problem can no longer be reduced to classifying an instance into a specific class, the instance model is expected to fail in this scenario, and our results are consistent with this.

When it comes to somatic mutations we expect some problems to require the model to learn the amount of a type of variant, and as a result labelled bags with two different specific amounts (Supplemental Figure 3C). We expected aggregations involving a sum to outperform aggregations involving a mean, but unexpectedly the instance sum aggregation did not solve the problem. Upon reflection, the output of the model immediately goes into the softmax of the crossentropy loss function, and as a result for the instance model and classification the distinction between a sum and mean is somewhat blurred. The fact that the sample model with a mean cannot solve this problem highlights a potential issue with using mean.

We can also envision problems with somatic mutations that require the model to be able to recognise the fraction of a type of variant, so we next labelled bags with two different fractions of an instance. As a first thought it might seem that only the mean aggregation will be able to solve this problem, but the sum contains all of the same information as the mean, and just requires further operations to convert the sum to the mean (Supplemental Figure 3D).

#### Regression Experiments

In addition classification tasks we can envision regression tasks that involve somatic mutations. To convert our model from one that solves a classification task to one that solves a regression problem all we have to change is the loss function (from binary crossentropy to mean squared error or negative partial likelihood for standard regression and Cox regression respectively). We only tested the sample models as they were the only models successful in the classification experiments. In Supplemental Figure 4A we had the bag label be a linear function of key instance count, whereas in Supplemental Figure 4B where the bag label was a quadratic function of key instance count. We also did a regression where the bag label depended on the fraction of bag filled with a key instance, and both the sample means and sums were able to the solve the problem, mirroring what we saw with classification (Supplemental Figure 4C).

#### Incorporating Sample Information

When it comes to including sample information into the model there are two obvious locations to do this: either immediately before the aggregation or immediately after. If added before, the attention will now be a function of instance and sample features. To explore what this might look like we performed some experiments with sample information. As seen in Supplemental Figure 5, there wasn't a clear performance difference whether the sample information was incorporated before or after the aggregation. The attention in Supplemental Figure 5A is what we might hope to expect from including

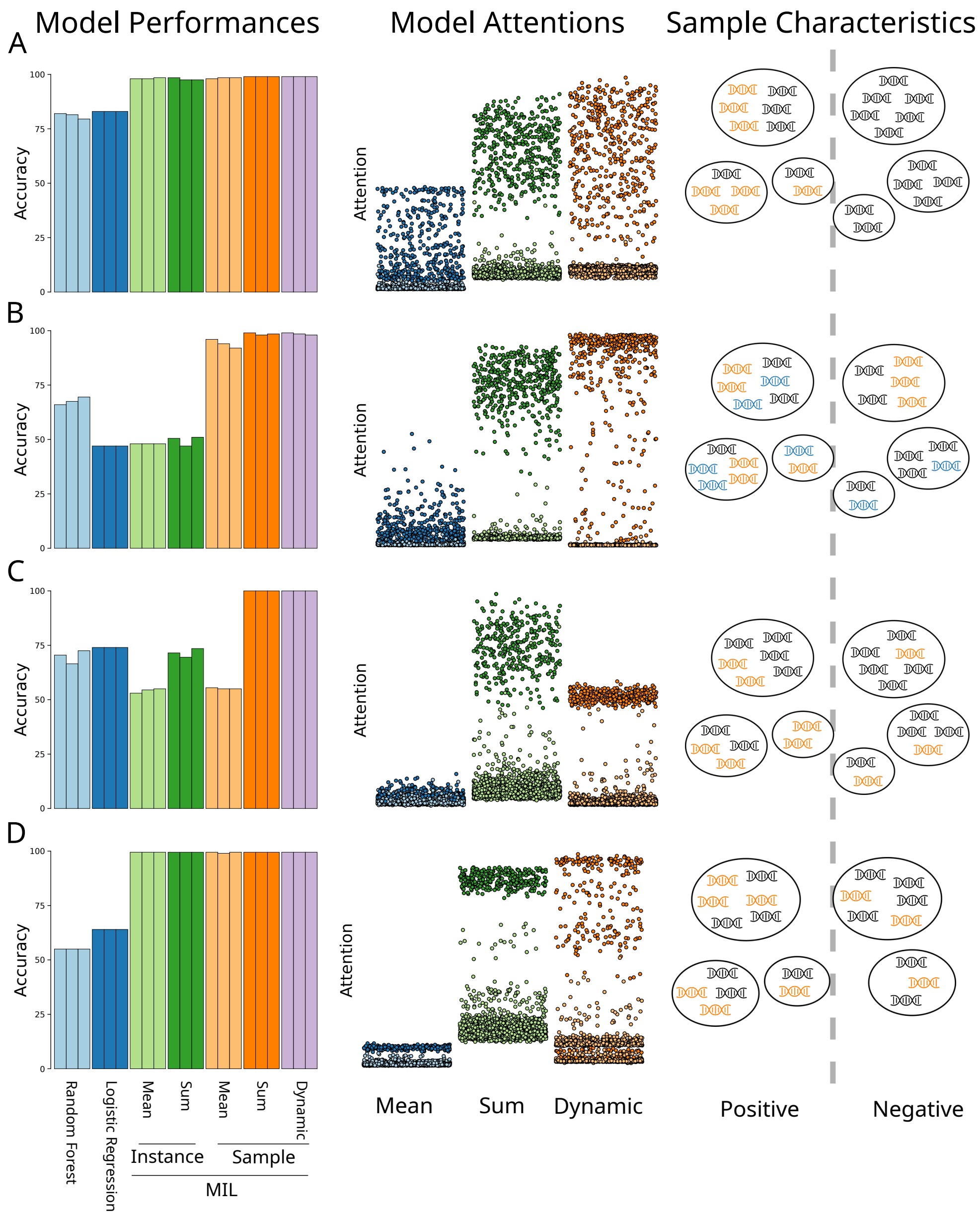


**Supplemental Figure 3. Sample-level classification with simulated data.** (A) Positive bags were labelled with a random fraction of a key instance, with negative bags having random variants. (B) A random fraction of each positive bag was filled with two key instances in equal proportions, and negative bags were composed of a random fraction of one of the two instances. (C) Positive bags were given a set amount of key instance regardless of bag size, and negative instances were given a different amount of key instance regardless of bag size. (D) Positive bags were labelled with a specific fraction of key instance, while negative bags were labelled with a different fraction. Key instances plotted as a darker shade in the attention figures. The evaluations and attentions are based on the test samples. Three runs were performed per model with the exact same train/valid/test split.

the sample information before the aggregation. In this experiment the key instances contribute to the bag value differently depending on the sample type, and this is correctly reflected in the attention. However, in Supplemental Figure 5B the sample information appears to be impacting both the value of the key instances and background instances, indicating that interpreting the attention when it is a function of instance features and sample features may not always be straightforward. Although in these examples there wasn't a striking performance difference, there were only three sample types. If the sample information is high dimensional then the instance-sample space will be very sparse, and how the sample information is included in the model may then impact performance.

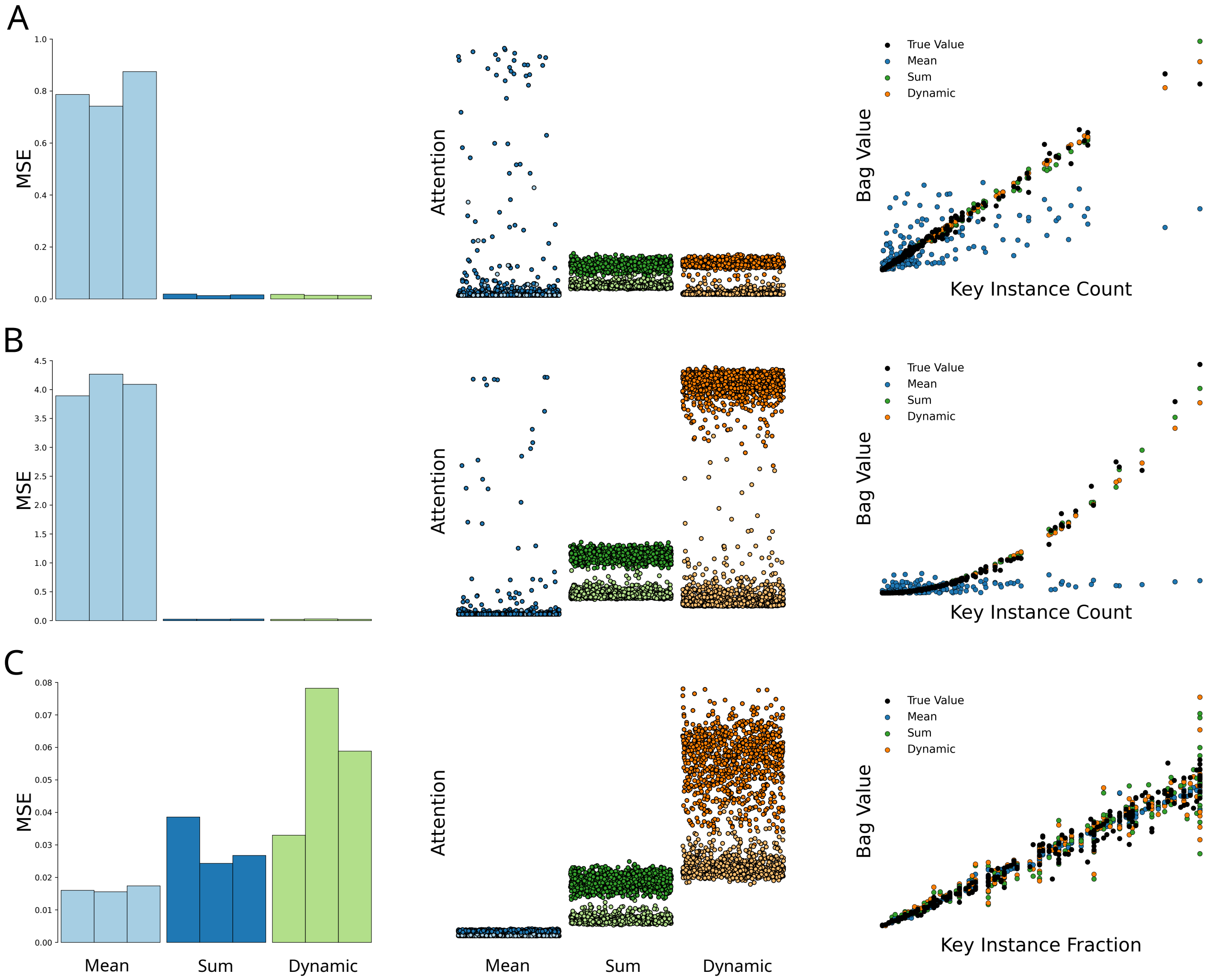


**Supplemental Figure 4. Sample-level regression with simulated data.** (A) Bag values were a linear function of the amount of key instance in the bags. (B) Bag values were a quadratic function of the amount of key instance in the bags. (C) Bag values were a linear function of the fraction of the bag occupied by key instance. In all cases some noise was added to the data. Key instances plotted as a darker shade in the attention figures. Three runs were performed per model with the exact same train/valid/test split.

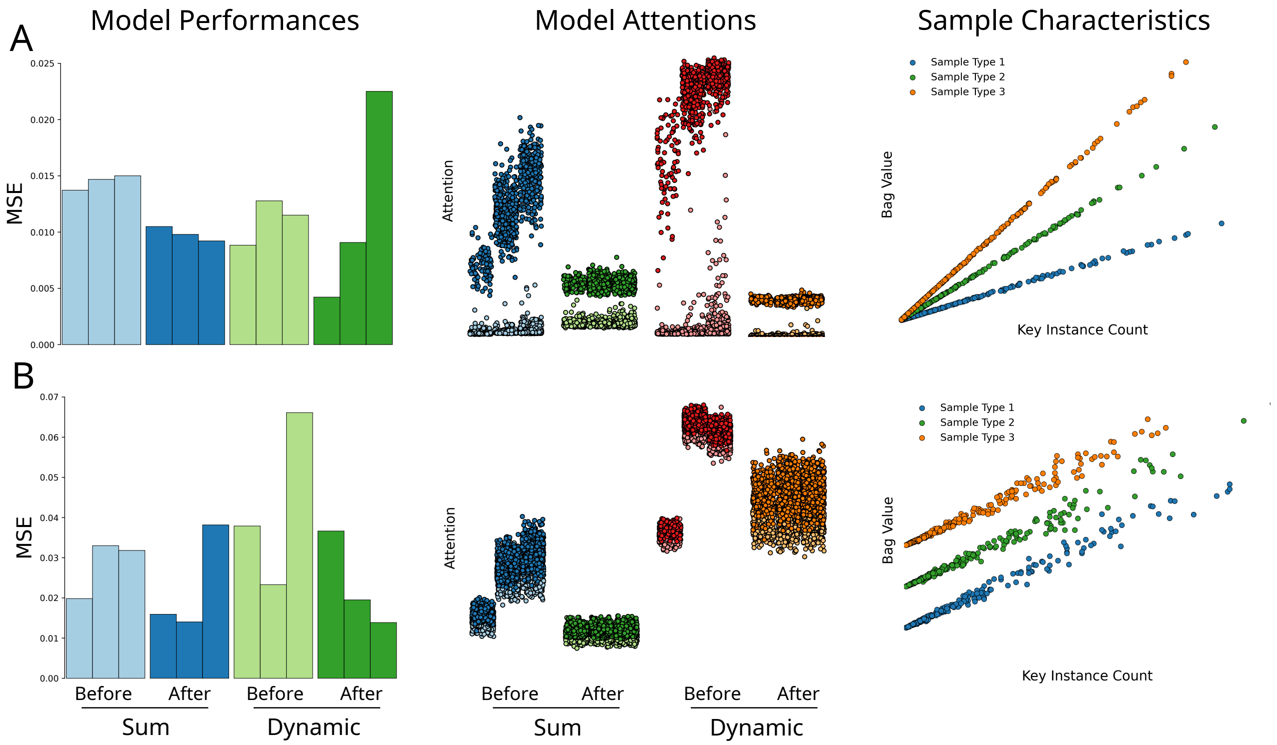


**Supplemental Figure 5. Incorporating sample information with simulated data.** Both experiments are regression tests where the bag value is determined by the key instance count in a linear manner, but in (A) the sample type affects how much each key instance contributes to the bag value, while in (B) it does not, but the bag value is affected by the sample type. Evaluations and attentions are based on the test samples. In the plot of attention the three columns per model are the three sample types, and key instances are plotted as a darker colour. Three runs were performed per model with the exact same train/valid/test split.

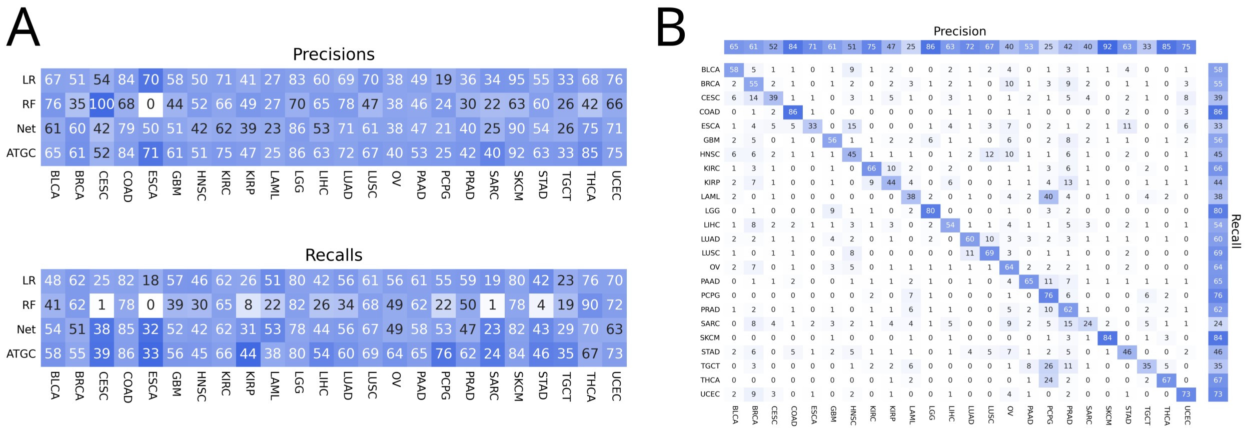


**Supplemental Figure 6. Tumour classification metrics.** (A) Precisions and recalls for the four models and gene as input. (B) Confusion matrix for ATGC and gene input.

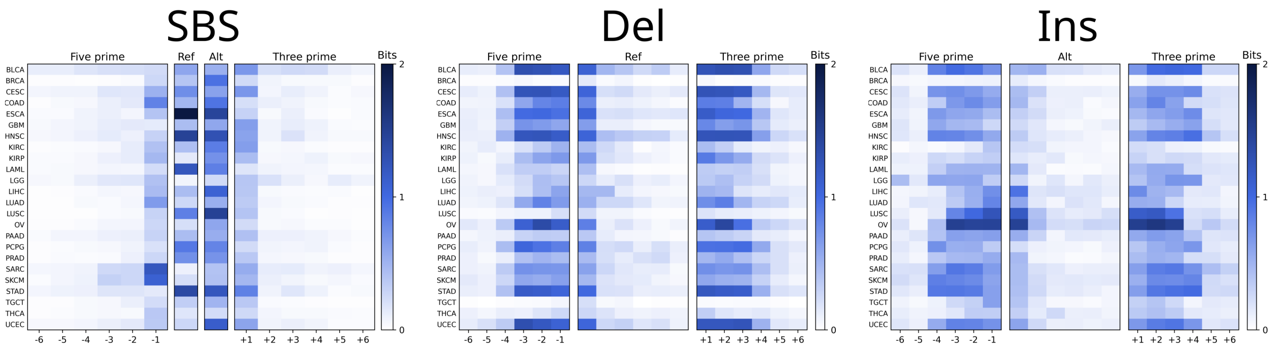


**Supplemental Figure 7. Information content of instance features.** The bits of information from sequence logos for the high-attention instances (top 5%) across all 24 attention heads for all instances in a single test fold.

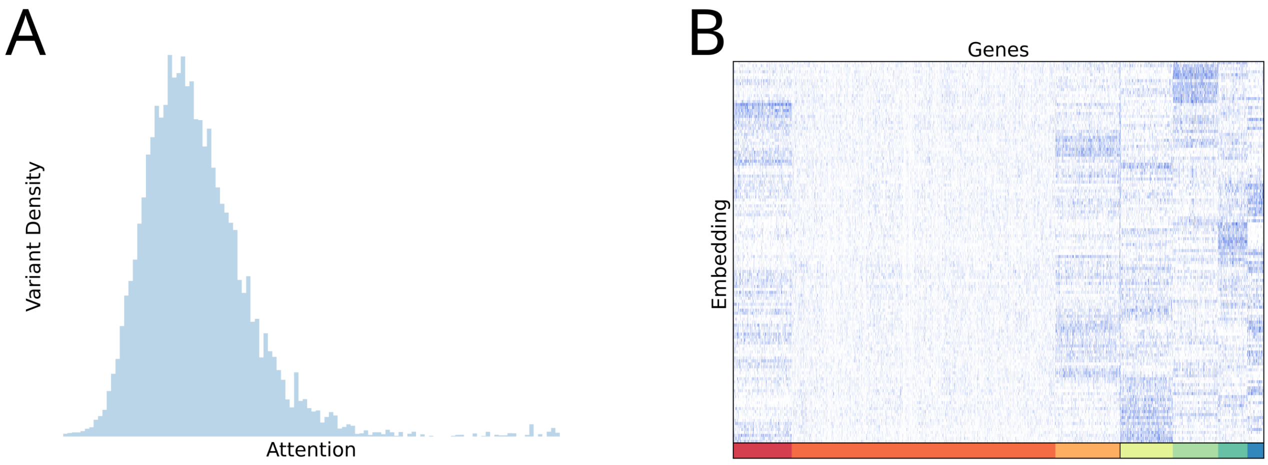


**Supplemental Figure 8. Gene attention and embedding matrix.** (A) Distribution of gene attentions for the SKCM head. (B) K-Means clustered gene embedding matrix. The genes in the black cluster are APC, ARHGAP35, ARID1A, ATRX, BRAF, CDH1, CDKN2A, CHD4, CIC, CTCF, CTNNB1, EGFR, FBXW7, FOXA2, FUBP1, GATA3, IDH1, KRAS, MAP3K1, NOTCH1, NRAS, PBRM1, PIK3CA, PIK3R1, PPP2R1A, PTEN, SMAD4, SOX17, SPOP, TP53, VHL.

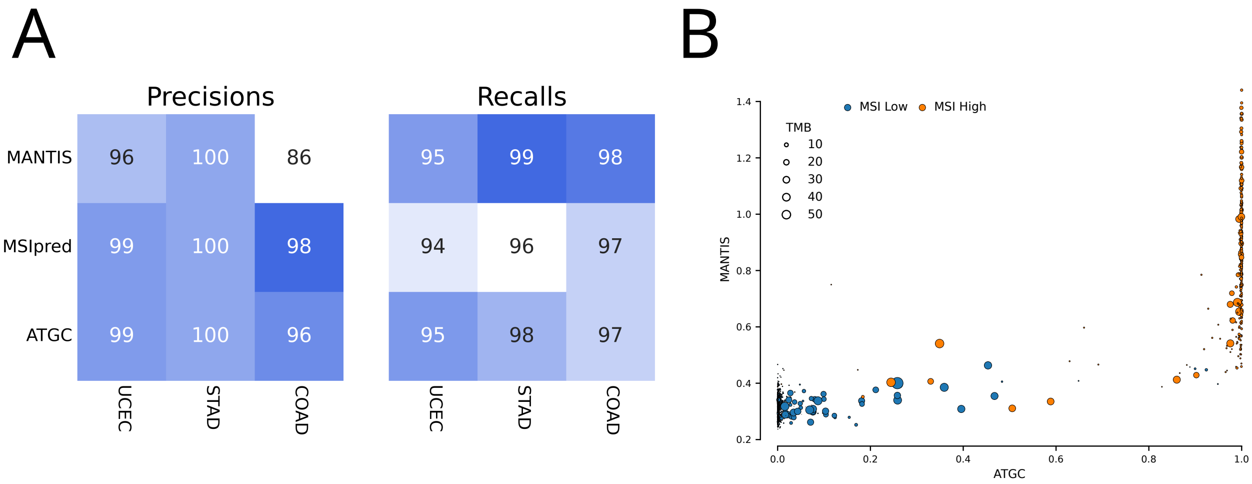


**Supplemental Figure 9. MSI predictions across cancer types and between models.** (A) Per cancer precisions and recalls for the 3 different models for the 3 most abundant cancer types. (B) MANTIS scores plotted against ATGC output probability showing high concordance of ATGC model output to MANTIS scores. Samples are colour coded by the PCR-based MSI status label, and the size of each sample corresponds to its total mutational burden (in thousands).

| Model | Accuracy | Weighted Accuracy | AUC |
|---|---|---|---|
| Logistic Regression | 81.2% | 77.8% | .977 |
| Random Forest | 72.7% | 66.1% | .960 |
| Neural Net | 80.8% | 78.3% | .980 |
| **ATGC** | **86.3%** | **83.3%** | **.986** |

**Supplemental Table 1. Tumour classification performance metrics for WGS and 96 context input.** Every model was trained with the same sample weighting and 10-fold cross validation.

| Data | Encoding | Aggregation | Model | Accuracy | Weighted Accuracy | AUC |
|---|---|---|---|---|---|---|
| 96 Contexts | Onehot | Sum | Logistic Regression | 44.4% | 42.5% | .916 |
| | | | Random Forest | 48.3% | 42.5% | .917 |
| | | | Neural Net | 47.6% | 45.8% | .928 |
| | | Weighted Sum | ATGC | 49.2% | 47.3% | .933 |
| 6 bp windows | Sequence Encoder | Weighted Sum | **ATGC** | **52.2%** | **50.7%** | **.940** |
| 1 Mb bins | Onehot | Sum | Logistic Regression | 46.6% | 42.2% | .898 |
| | | | Random Forest | 43.8% | 39.2% | .880 |
| | | | Neural Net | 49.6% | 46.6% | .918 |
| | Embedding | Weighted Sum | **ATGC** | **52.3%** | **49.0%** | **.924** |
| 30 kb bins | Onehot | Sum | Logistic Regression | 52.2% | 47.8% | .925 |
| | | | Random Forest | 46.1% | 41.5% | .887 |
| | | | Neural Net | NA | NA | NA |
| | Embedding | Weighted Sum | **ATGC** | **54.7%** | **50.6%** | **.928** |
| Gene | Onehot | Sum | Logistic Regression | 55.3% | 50.8% | .932 |
| | | | Random Forest | 48.1% | 44.4% | .894 |
| | | | Neural Net | 53.8% | 51.1% | .932 |
| | Embedding | Weighted Sum | **ATGC** | **58.6%** | **54.7%** | **.940** |

**Supplemental Table 2.** **Tumour classification performance metrics for exome NCIt codes.** Every model was trained with the same sample weighting and 5-fold cross validation. When using a large input vector like the 30 kb bins our procedure for optimising neural nets cannot be used.
